# Flock response to sustained asynchronous predator attacks

**DOI:** 10.1101/2023.11.14.567144

**Authors:** Siddhant Mohapatra, Pallab Sinha Mahapatra

**Affiliations:** Department of Mechanical Engineering, Indian Institute of Technology Madras, Chennai, India 600036

## Abstract

Collective behaviour is a ubiquitous emergent phenomenon where organisms share information and conduct complicated manoeuvres as a group. Dilution of predation risk is presumed to be a major proponent contributing towards the emergence of such fascinating behaviour. However, the role of multiple sources of predation risk in determining the characteristics of the escape manoeuvres remains largely unexplored. The current work aims to address this paucity by examining the response of a flock to multiple persistently pursuing predators, using an agent-based approach employing a force-based model. Collective features such as herding, avoiding and split-and-join are observed across a wide spectrum of systemic conditions. The transition from one response state to another is examined as a function of the relative angle of predator attack, a parameter exclusive to multi-predator systems. Other concomitant parameters, such as the frequency of attacks and compatibility of target selection tactics of the predators, have a significant effect on the escape probability of the prey (i.e., the success rate of escape manoeuvres). A quantitative analysis has been carried out to determine the most successful combination of target selection while also focusing on beneficial ancillary effects such as flock splitting. The long-term dynamics of the system indicate a faster decay of prey numbers (higher prey mortality) at higher coordination strength due to a monotonically decreasing relation between coordination strength and prey speed supplanted by coincidental synchrony of predator attacks. The work highlights the non-additive nature of the effects of predation in a multi-predator system and urges further scrutiny of group hunting dynamics in such systems.

**Author summary:** Collective motion is a natural phenomenon observed across a wide range of length and time scales. One purported reason for the development of such behaviour is to reduce the individual risk of predation through the many-eyes effect and group manoeuvring in case of attacks. However, the behaviour of the prey flock can turn out to be starkly different when there are multiple predators involved. We examine the response of the flock in the presence of multiple predators and find the circumstances leading to the occurrence of different escape manoeuvres. We observe the stricter penalty warranted on the flock due to certain manoeuvres, such as split-and-join, due to the asynchronous and persistent nature of the predator attacks. We also identify the issues with superfluous coordination among prey and its ramifications in terms of prey mortality. The combined effect of the predators is found to outpace the sum of individual predator prowess. The current work emphasises the distinct dynamics of a multi-predator system and puts forth pertinent queries regarding synchronisation among predators and group hunting tactics.

## Introduction

Collective behaviour is an exquisite phenomenon observed across multiple scales in nature. From an evolutionary standpoint, the risk of predation remains a plausible explanation for the origin and development of such phenomena. Works such as that by Ioannou et al. [1] showcasing natural predators preferentially attacking isolated over aggregated virtual prey and Olson et al. [2] presenting swarming as a consequence of predator confusion highlight that predation risk is sufficient for the evolution of collective motion. Collective behaviour has been reported to reduce the individual burden of vigilance against predators and dilute the predation risk apart from causing predation confusion [3–5]. The early detection of predators’ approach and fast information transfer due to flocking allow the prey to perform well-coordinated escape manoeuvres. Magurran & Pitcher [6] reported a wide array of evasive manoeuvres undertaken by minnows (*Phoxinus phoxinus*) and the frequency of their occurrence in response to pike (*Esox lucius*) attacks. Compact, inspect and avoid were among the most commonly observed manoeuvres. Similarly, Storms et al. [7] reported the escape manoeuvres, such as blackening, wave event, split, and merge, to name a few, in a starling (*Sturnus vulgaris*) flock under threat of predator from a raptor. They examined the circumstances surrounding the incidence of each evasive measure and connected them to the threat intensity of the falcon (including the speed of the attack) and the state of the flock before the attack. Papadopoulou et al. [8] analysed trajectories of pigeon flocks chased by a robotic falcon and developed a model to demonstrate the relationship between the propensity for turning and distance from the predator, leading to collective turn negotiation. Domenici and Batty [9] discussed how schooling helps herring (*Clupea harengus*) to rectify their trajectories while turning in response to an acoustic disturbance. Palmer and Packer [10] investigated the response of wild impala, zebras and wildebeest to hi-def life-sized images of common African predators. They reported that the nature and intensity of evasive manoeuvre varies for each species and is a function of the predator’s hunting style and the prey’s risk perception.

Taking a broader overview of animal behaviour-based studies, a fair amount of literature is concerned with observations in the wild [10–13] as well as experiments in controlled environments [14–17]. Such hands-on approaches help establish a broad relation between the characteristics and the behaviour of the organisms. However, isolating a causal link for specific behavioural traits becomes increasingly difficult. At the same time, the ethical and other procedural clearances associated with experimentation on macro-scale animals supplant the arduousness. Computational models play an important role in uncovering the underlying mechanisms of social behaviour in such cases. Agent-based approaches, in particular, have been a promising technique for simulating collective behaviour in organisms, as they provide an adequate resolution (“collective” scale) for study while also allowing individual-level control. In the past few decades, researchers have developed numerous mathematical models such as the Couzin model [18], the Boids’ model [19], the Kuramoto class of models [20] among others, to define how organisms communicate with each other in order to maintain the structural integrity of the collective. Cellular automaton models have also been put to use to simulate prey motion and their interactions with predators as well as the environment [21–23]. Of late, evolutionary and genetic modelling [24, 25], models based on game theoretic approaches [26, 27], and models based on reinforcement learning and neural networks [28, 29] have also been pitched to varying degrees of accuracy. All of these models attempt to discern the information transfer within a flock, which leads to the concerted evasive manoeuvres observed in nature. An optimised information transfer network could be emulated in robotic swarms for far-reaching implications in the military and the disaster rescue sectors [30].

Interactions between adversarial species, aptly named predator-prey interactions, are a major talking point in the biophysics community due to the sheer complexity involved. A significant section of the spawned studies have focused on the transmission and role of visual, olfactory and auditory cues when predators and prey are in proximity [31–34]. Barring a handful of studies [35–40], the majority of the extant literature (esp. computational) is concerned with single predator-prey encounters rather than successive or concerted predator attacks. However, experimental observations have highlighted the ubiquity of group hunting in certain ecosystems [41–44]. The reviews by Sih et al. [45], Bailey et al. [46], and more recently Hansen et al. [47] accentuate the benefits of group hunting while emphasising the dearth of comprehension of the underlying physics involved in such events. The attack patterns of predators have been explored in the literature to some extent [10, 24, 48–50]; however, there is a lack of studies highlighting the effect of predator attacks using a combination of these strategies.

The current work focuses on explaining the aftermath of repeated predator attacks on a generic prey flock using an agent-based model. Emphasis has been placed on explaining the prey response to different attack strategies and the transitory and long-term behaviour of the system. The asynchrony and persistence of attacks by multiple predators are found to have a substantial impact on individual hunting success. From the prey’s perspective, the costs associated with alignment have been revisited and commented upon. The next section details the numerical model, while the subsequent section describes the qualitative behavioural response of the prey to the predator attacks, followed by a quantitative analysis.

## Numerical methodology

### Model

An agent-based modelling approach has been considered for the study, wherein all the entities are assumed to be agents placed in a periodic domain. An underdamped Langevin model with non-negligible agent inertia has been considered. In this work, the agents can be classified into two categories: a prey agent and a predator agent. There are physiological differences between the two species.

#### Prey agent

The governing equation for a prey agent *p* consists of a propulsion term, a short-range pairwise separation term, a mid-range alignment term and a long-range attraction term (see Eq. 1).

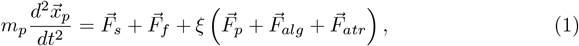

where, *m*_*p*_ and 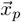 are the mass and the position of the prey agent respectively, 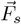 is the pairwise separation force, 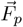 is the propulsion force, 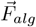 is the alignment force, 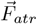 is the attraction force, 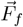 is the frictional force and *ξ* is the state of existence of the prey (i.e., *ξ* is 1 for live prey and 0 for dead prey).

The present work considers a zonal classification around the prey agent for the action of the forces. Every prey agent has a separation zone, an alignment zone, and an attraction zone, based on which the forces act on the prey agent (see Fig. 1). The separation force acts only within the limits of the separation zone (*r*_*sep*_), and the alignment force acts in the alignment zone sans the separation zone (*r*_*alg*_− *r*_*sep*_), while the attraction force acts in the area between the alignment zone limits and the attraction zone limits (*r*_*atr*_ − *r*_*alg*_). Apart from these zones, the prey agent has a detection zone encompassing the entire zonal area, where the agent can detect the presence of the predator. Assuming the sensory input to be predominantly visual [31], a blind angle *θ*_*b*_ is considered symmetrically about the direction opposite to the motion of the agent.

**Fig 1.**
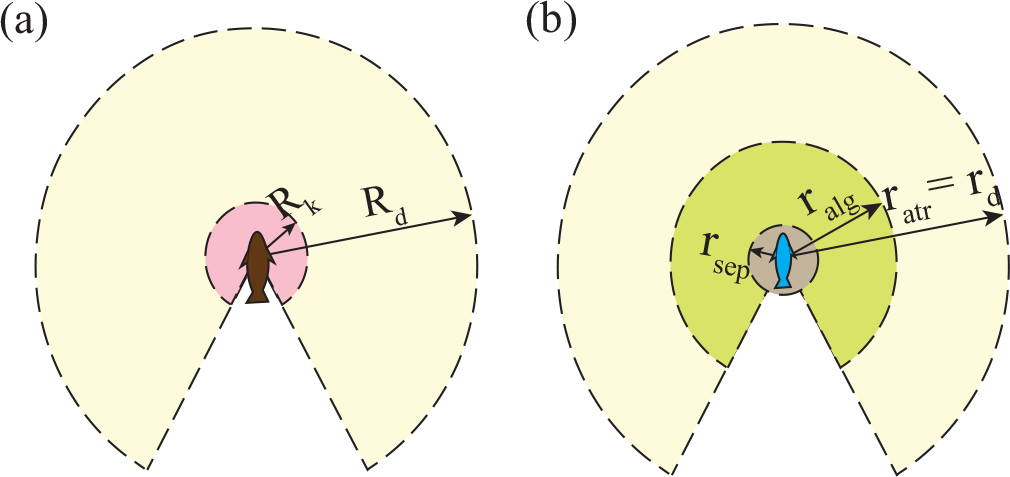
The zonal structures of (a) the predator and (b) the prey agents are elucidated; Notations for the predator are k: kill, d: detection; while for the prey, sep: separation, alg: alignment, atr: attraction, and d: detection zones respectively.

The pairwise separation force is a short-range repulsive force that prevents the overlap of agents. The force acting to separate any two agents *i* and *j* can be represented as Eq. 2

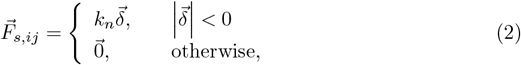

where, 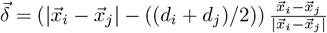, is the separation between the two agents (in vectorial form) and *d*_*i*_ and *d*_*j*_ are the diameters of the respective agents.

The propulsion force begets the term “active matter” and is the force responsible for the movement of the agents in a desired direction. It could, otherwise, be defined as the will of the agent and is accountable for the non-equilibrium nature of active systems. In the present work, the prey agent has two states of activity: cruising state and alarmed state (see Eq. 3). The cruising state occurs in the absence of any danger (predator) in the vicinity, and the line of action of the force is in the direction of the current velocity. A prey in an alarmed state tries to move away from the predator in its neighbourhood as fast as possible; hence, the line of action of the propulsion force is in the direction away from the predator’s position.

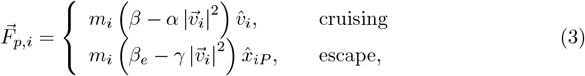

where, 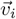 is the velocity of the prey agent 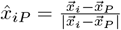 is the direction of the shortest approach from the predator *P* towards the agent *i, β* and *β*_*e*_ are the cruising and escape accelerations respectively and *α* and *γ* are the Rayleigh friction coefficients to prevent unbounded accelerations.

The alignment force is the cause of the flocking phenomena and is a force-based interpretation of Vicsek’s alignment rule [51], where every agent tries to align its velocity to that of its neighbours (see Eq. 4). A metric distance-based model is considered for the neighbourhood definition, as opposed to the kNN or the Voronoi neighbours approach.

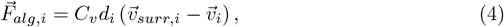

where, 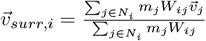 is the weighted average velocity of the live prey neighbours (*N*_*i*_) of agent *i* (a Gaussian weighting function centred at *i* and depending on distance 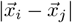 is considered), *d*_*i*_is the characteristic dimension of the agent (here, the diameter), and *C*_*v*_ is the strength of alignment among the prey. An orientational white noise *ζ* with ⟨*ζ*(*t*)*ζ*(*t*^*′*^)⟩ = *σ*_*ζ*_*δ*_*ij*_*δ*(*t* − *t*^*′*^) is superposed on the direction of the alignment force, similar to [52].

In cases where a certain prey gets separated from its flock, the long-range attraction force acts to drive the isolated prey towards a nearby flock. The form of the attraction force (initially defined by Warburton & Lazarus [53]) is such that the farther an isolated prey perceives the nearest flock to be, the more vigorous is the drive to cover the distance (see Eq. 5). As the prey gets closer to the flock, the attraction force eases off.

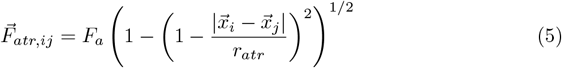

where, *F*_*a*_ is the strength of the attraction force and *r*_*atr*_ is the radius of the attraction zone.

The frictional force 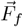, as the name suggests, acts opposite to the direction of motion of the agent. The frictional drag experienced by the organism due to the relative velocity with respect to the surrounding medium can be approximated, following Viscido et al. [54] (see Eq. 6).

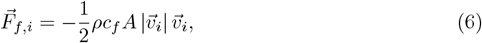

where *c*_*f*_ is the drag coefficient, *A* is the surface area in contact with the fluid, and *ρ* is the density of the surrounding medium.

#### Predator agent

The governing equation for a predator agent *P* consists of a propulsion term and a short-range pairwise separation term (see Eq. 7).

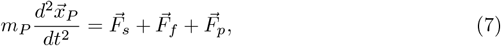

where the RHS represents the sum of the pairwise separation force 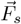, the frictional force 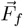 and the propulsion force 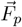. *m*_*P*_ and 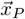 are the mass and the position of the predator agent, respectively. The current work assumes the predator agent to be alive throughout the simulation. The zonal classification for the predator constitutes a kill zone *R*_*k*_ and a detection zone *R*_*d*_ (see Fig. 1). Any target prey which enters the kill zone is killed, while the predator can scan the live prey in the detection zone before selecting a target.

The pairwise separation and the frictional forces for the predator agent have the same form as in the case of prey agents. However, the differences lie in the case of the propulsion force. The predator agent has three states of activity: satisfied state, refocus state and pursuit state. The satisfied state occurs when the predator is successful in hunting the previous target prey, while the refocus state occurs in case of failure of the same. The propulsion force, in both the aforementioned states, acts in the direction of the current velocity, similar to that of the prey, while in the pursuit state, the predator propels itself in the direction of the shortest distance towards the target prey (see Eq. 8).

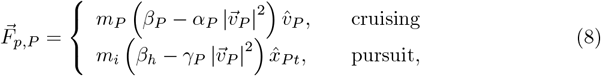

where *β*_*P*_ and *β*_*h*_ are the cruising and the hunting accelerations of the predator, *α*_*P*_ and *γ*_*P*_ are the Rayleigh’s friction coefficients to prevent unbounded acceleration,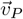 is the velocity of the predator and 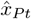 is the direction from the predator towards the target prey.

The predation process, as defined in one of our earlier works [55], consists of multiple steps: a) detection stage, b) target selection, c) pursuit stage, and d) satisfaction/refocus stage. The predator detects all live prey in its vicinity and selects a target from among them. The predator, then, pursues the target prey till it either captures it or the hunting time expires (whichever is earlier). The maximum hunting time ensures a limited duration for the predator to exert a high acceleration (after which it gets tired). If the predator is able to capture the prey, it experiences a satisfaction stage where it doesn’t pursue any prey, while a failed hunt results in the predator requiring some time to recuperate from the exertion (a refocus time). The details of the process have been illustrated in S1 Fig. In nature, the predator selects its target prey based on a number of environmental as well as internal factors. In the present work, we assume visual cues as the primary source of information and two possible ways of target selection: head-on attack and split attack. In the head-on attack (see Fig. 2(a)), the predator attacks the live prey nearest to its position, while in the split attack (see Fig. 2(b)), the predator attacks the live prey closest to the algebraic centre of the visible flock. The nomenclature follows the norm of the outcome, as a head-on attack leads to the pursuit of a single prey, while a split attack often causes the division of a large flock into smaller divisions.

**Fig 2.**
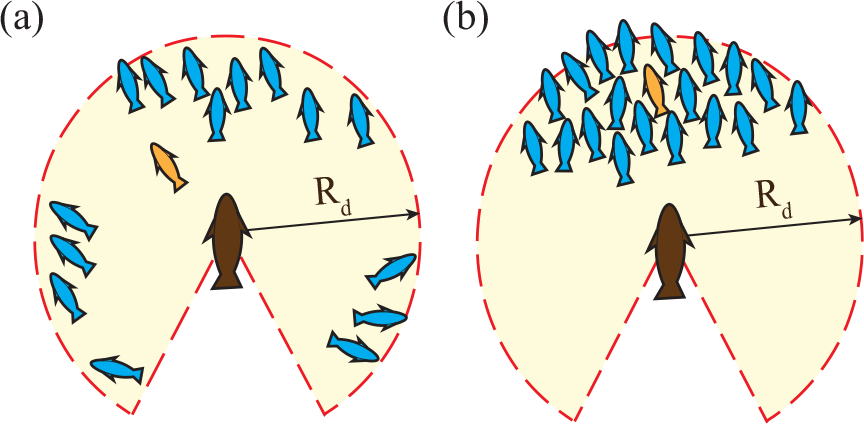
The different strategies that the predator can adopt while attacking prey flocks are showcased here: (a) head-on attack and (b) split attack. (Note: The blue prey are possible targets, the orange prey is the selected target prey, and the predator is coloured brown.)

### Simulation details

The system under observation constitutes a flock of prey and two predators in a square periodic domain of side *L*. Both the prey and the predator agents are assumed to be soft disks and can occupy off-lattice positions. The predators kill the prey; however, they consume a minuscule part of it such that there is no difference in the predator mass. The forces acting on each of the agents are calculated at every time step, and the velocities and displacements are updated synchronously using the Velocity-Verlet algorithm. The present work focuses on predation effectiveness and the prey response to a combination attack by the predator duo. As a consequence, different cases have been considered with different permutations of the two hunting strategies described in Sec. Predator agent. In the case of the combination attack, every predator has an equal probability of choosing a head-on or a split attack for each encounter. The detailed list of parameters and their values considered are tabulated in S1 Table. Fifty realisations have been run with different starting positions of the predators with respect to the prey flock for each of the parametric cases. The simulation has been carried out using an in-house C++ code, and the data has been processed and plots generated using MATLAB and *seaborn* package in Python 3.8.

## Results

The current work focuses on the behavioural response of a flock of prey to multiple predators. A qualitative response analysis has been carried out to determine the precursors leading up to certain escape manoeuvres demonstrated by the prey, This is followed by a quantitative take on the long-term escape success dynamics of the prey, taking into account parameters such as agreement among predators’ target selection strategies and the impact of persistence and asynchrony (and coincidental synchrony) of predator attacks.

### Prey response

Across the simulation time, the prey is observed to perform large-scale coordinated manoeuvres to escape the predator, similar to those found in nature. The occurrence of such manoeuvres depends largely on the flock’s heading and the predators’ relative approach. The collective manoeuvres observed among the prey are elucidated in Fig. 3. The first event is termed the “split” manoeuvre and occurs when the predators attack a flock from opposite sides (see Fig. 3(a)). In Fig. 3(a), the predators are denoted by *P*_1_ and *P*_2_, while the prey flocks are denoted by their centre of mass *F*_*i*_ (where *i* is the flock designation). The path traced by the two predators *P*_1_ and *P*_2_ shows the approach from opposite directions towards the undivided flock *F*_0_ (also see Fig. 3(a-I)). The time progression of the event is indicated through the colour of the markers, where 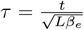 is the non-dimensional time of the event across which the event occurs. As the predators move closer to the flock *F*_0_, the prey attempt to move in a direction away from both the predators. Based on the proximity to the respective predators, the flock’s attempt to escape causes a constriction at the location nearest to both the predators (see Fig. 3(a-II)). The constriction continues narrowing, and the initial flock (*F*_0_) eventually splits into smaller divisions (*F*_1_ and *F*_2_), which start moving in opposite directions (as depicted from the colour profile), and the predators follow the flock in their immediate neighbourhood (see Fig. 3(a-III). The second event is termed the “herding” manoeuvre, where the prey performs a directed motion away from the predators’ approach. The uni-directional movement of the prey is a consequence of the predators attacking from similar angles (see 3(b)). The third class of events is referred to as “individual attack”, wherein a predator chases a particular flock exclusively in the presence of multiple flocks in the domain. Such pursuits are equivalent to two separate systems with a single predator chasing the flock (see Fig. 3(c)) and usually follow a split manoeuvre. The relative angle of attack of the predators plays a major role in determining the prey response. Figure 3(d) shows the departure from a herding response at lower relative attack angles (Ω_*P*_) to a mostly split response from the flock as Ω_*P*_ increases. However, at intermediate values of Ω_*P*_, the flock can respond either by herding or by splitting (as shown in the shaded region in Fig. 3(d)). While it is relatively straight-forward that the attack direction of the predators has a clear bearing towards the prey response, the coordination among the prey also serves as a proponent to prefer splitting over herding, especially at intermediate Ω_*P*_ (see Fig. 3(d) inset). A possible reason for the preference is the localised nature of the alignment interactions (the alignment kernel decays exponentially as the distance from the agent increases).

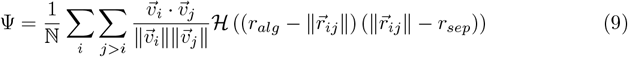

where, 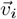 is the velocity of the agent *i*, ℕ is the number of unique neighbour pairs among the live agents, and ℋ is the Heaviside function.

**Fig 3.**
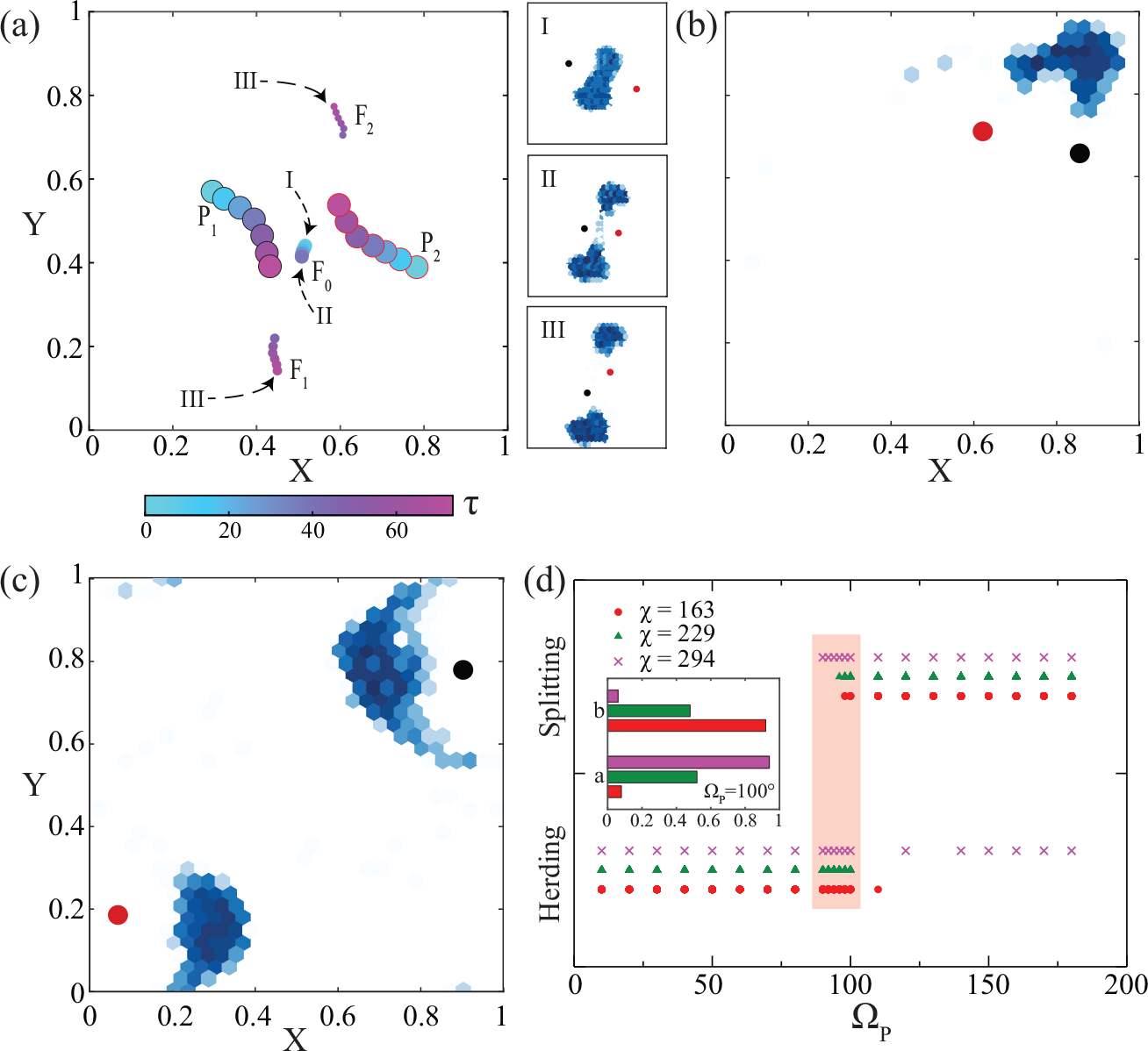
The majority of the events occurring in the system fall into either of the three categories: case (a) where the two predators (*P*_1_ and *P*_2_) attack a flock (*F*_0_) from opposite directions, causing the flock to split into smaller flocks *F*_1_ and *F*_2_ (the flocks are represented by their centre of mass and the colour bar shows the time progression of the event, where 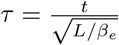 is the non-dimensional time), case (b) where the two predators attack a flock from a similar direction, leading to a herding scenario, and case (c) where each of the predators attacks different flocks, effectively reducing the system into two single predator-prey systems. In case (a), the predator positions are tracked alongside the centroids of the flocks before and after division. Panels (a-I) through (a-III) depict the flock division phenomena in the form of time-lapse prey density plots. Cases (b) and (c) elucidate the prey density plots along with the predator positions. Panel (d) depicts the transition from the splitting (case a) to the herding regime (case b) with the increase in the relative angle of attack of the predators Ω_*P*_ for both predators following head-on attack and with different degrees of prey coordination 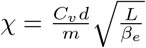. The inset depicts the transition from a majorly herding case to a majorly splitting case as the coordination among the prey increases for Ω_*P*_ = 100° (similar behaviour is observed throughout the shaded region). (Note: This image serves as a representative of the numerous events occurring in the system across time.)

The events showcase different manoeuvres such as split and join (Fig. 3(a)), herd (Fig. 3(b)), and avoid (Fig. 3(c)), which are also widely observed in natural flocks [6, 56]. The split-and-join phenomenon is particularly interesting in the scope of this work as it highlights the effect of multiple attacking predators. Figure 4 demonstrates the different stages of the split-and-join phenomena observed in the system at lower coordination values: a polarised stage (shaded blue), a confusion stage (shaded red) and a recovery and splitting stage (shaded green). In the first stage, two highly polarised flocks (Ψ ≈ 0.96) moving with an average speed typical of *χ* = 163 are put on a path of imminent collision due to persistent pursuit by their respective predators (refer Fig. 4 inset (a)). As the two flocks collide, it leads to a temporary disruption of order (confusion), indicated by a sharp reduction in the local order parameter Ψ (refer to the agent orientations in Fig. 4 inset (b)). The average speed of the united flock 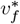 is also negatively affected due to the head-on collision. However, due to the alignment interactions between the agents (the strength of which is quantified by *χ*), the polar order is eventually recovered in the system (Ψ starts increasing again; refer to the ordering of agent orientations from inset (c) to inset (d)). Meanwhile, the two predators approach the united flock from opposite directions, compelling the agents to move in a direction transverse to the line of the predator attack (see Fig. 4 inset (e)) and leading to the formation of a constriction as explained a priori in Fig. 3(a). The dual predation pressure causes an increase in the prey speed in the transverse direction as the constriction becomes sparser (see inset (f)), eventually culminating in a split of the united flock into two highly polarised flocks (inset (g)). The split-and-join manoeuvre, observed in Fig. 4 has a unique signature of a transient decrease in local order, followed by an increase in flock speed. The absence of either of the changes points to disparate scenarios. Split-and-join manoeuvre has been observed, even in the presence of a single predator [6]; however, in this particular variation, due to the multiple predators, the time associated with the confusion stage is a hefty disadvantage for the prey (due to slower response times and reduced coordinated evasion capabilities). The persistent predator attacks also result in a highly transitory conjoined state of the flocks, thereby preventing the subsistence of large flocks.

**Fig 4.**
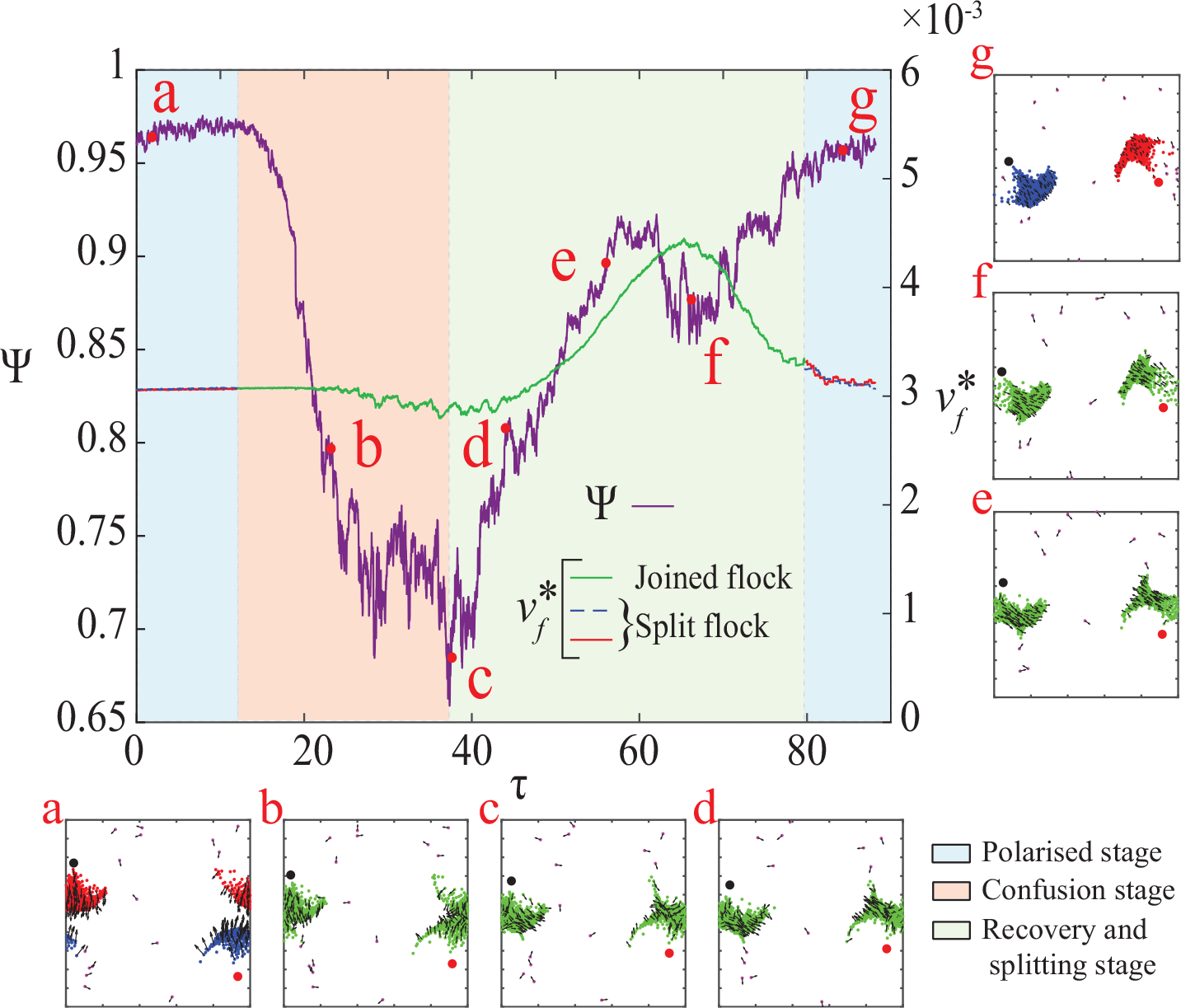
The local order parameter Ψ and the non-dimensional average speed of the flock 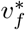are plotted against time *τ* for the duration of the split-and-join manoeuvre with snapshots of the system inset. In the first stage (shaded blue; polarised state), there are two distinct polarised flocks (as shown in inset (a)) moving in a path for imminent collision. Once the collision of the two flocking fronts occurs, there is a transient reduction in the local order (shaded red; the confusion stage). The insets (b) and (c) show the progressively dissimilar orientations of the colliding flocks. The flock, however, recovers from the confusion (indicated by recovering Ψ; shaded green) due to the alignment interaction (inset (d)) until opposing predation pressures cause stretching of the united flock (indicated by a rise in 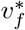 from normal levels) in a direction transverse to the line of attack (inset (e) and (f)). The narrowing of the constriction eventually leads to the splitting of the flocks, which now exhibit polarised motion, thereby completing the cycle (inset (g)). (Note: The red and the blue agents in the insets correspond to the two polarised flocks, respectively, while the green particles represent a joined flock.)

### Quantitative analysis and hunt statistics

The previous section gives an overview of the various events concerning predators’ attacks and the subsequent prey responses. However, there is also a need to understand the changes in the prey response with respect to the environmental conditions and to assess the efficiency of the predator in hunting down the prey under such circumstances. The changes in hunting efficiency can be attributed not only to the predator’s different target selection strategies but also to the disposition of the prey. The predator can opt for a head-on or a split attack (refer Sec. Predator agent). On the other hand, there are multiple methods to implement the herding propensity of the prey numerically. One approach considered by Chakraborty et al. [57] is to change the interaction radius of the prey, which effectively alters the number of aligning neighbours. However, interpreting the interaction range for an organism as a time-varying function seems unlikely in the absence of external mitigators of sensory input. In the current work, the degree of the willingness to herd has been represented through a coordination parameter *χ*, where a higher value of *χ* leads to a stronger herding tendency. However, a stronger herding doesn’t necessarily translate to a successful evasion when there are multiple predators involved. To determine the effect of a higher *χ*, a scaling analysis has been carried out considering all the relevant forces, and the speed has a quadratic decay with the coordination parameter *χ*, ceteris paribus. Assuming a well-distributed but compact flock, the pairwise separation and the attraction forces can be ignored, and the force balance for prey in an alert state is delineated in Eq. 10.

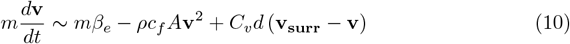

The scaling analysis is considered over a small time window (such that 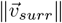 can be assumed constant), leading to Eq.10 reducing to a quadratic relation (see Eq. 11) for the parameter space considered in the simulations (refer S1 Table). The detailed derivation of the scaling analysis of the speed of the prey can be found in the S1 Appendix.

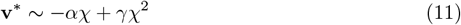

where, 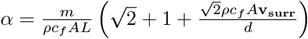 and 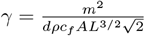 are positive constants and 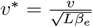 is the non-dimensional speed of the prey. The reduction of speed of the prey at higher coordination, indicated in the scaling relation, corroborates with that observed in the simulations (see Fig. 5) and can be attributed to the increased cost of maintaining the velocity alignment. As a result, however, the overall speed of the flock reduces, making it more vulnerable to subsequent predator attacks. The presence of multiple predators exacerbates the situation as the predators attack asynchronously and can lead to higher killing in case of consecutive attacks. The asynchrony and coincidental synchrony between the predators is revisited in S2 Fig.

**Fig 5.**
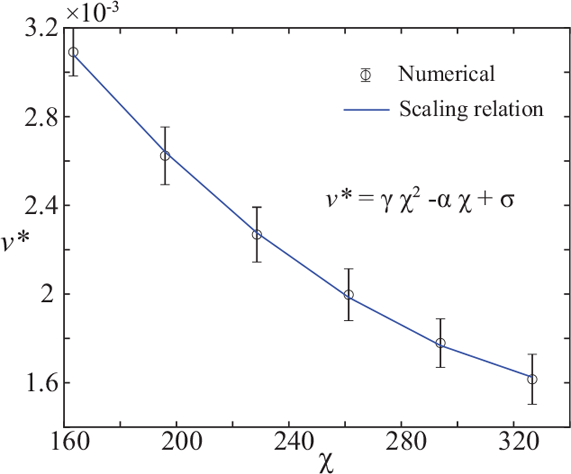
The correlation between the speed of the prey and the coordination coefficient *χ* is illustrated and is in good agreement with the quadratic relations obtained from scaling analysis. (Note: 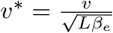 is the non-dimensional speed of the prey.)

The predators’ hunting prowess can also be studied by examining the rate at which it is able to kill prey. Figure 6 depicts the decay in live prey numbers due to predator hunting as time passes by, and the insets illustrate the rate of killing *α*_*K*_ of live prey with time at different degrees of coordination *χ*. To note, there is a limited number of prey in the system, and dead prey are not replaced with new live prey. As explained earlier, a head-on attack by the predators has a faster rate of killing prey compared to a combination attack (the rate is at least 20% higher for a head-on attack). The adverse effect of coordination, causing a slowdown in the prey movement (refer Fig. 5), can be seen by comparing the *N*_*l*_ decay curves for *χ* = 163, *χ* = 229 and *χ* = 294. The rate of killing is lower in the case of lower coordination, owing to higher speeds and greater freedom for individual movement. Regardless of the individual attack strategies of the predators, a linear trend of the rate of killing with time is observed for different prey coordination strengths *χ* (except *χ*163) for the majority of the simulation time. The gradual decline in the rate of killing *α*_*K*_ can be explained by a gradual reduction in the available prey numbers. It might also be an artefact of the predation model, which considers a limited hunting time, a refocus time and a satisfaction time (see Sec. Predator agent). The details of the temporal dependence of the number of live prey *N*_*l*_ and the rate of killing are analysed in S2 Appendix. When there is an abundance of prey options for the predators, the predators are able to kill the prey successfully within the stipulated hunting time, which is evidenced by an ever so slightly decreasing killing rate *α*_*K*_ (see Fig. 6 insets) for most of the *τ* values. However, when the live prey become very sparse in the system (at high *τ*), a rapid decrease in the rate of killing is observed (see insets of Fig. 6), as the dead prey occlude the vision of the predators and therefore, the predators have to search for a longer duration before encountering any live prey. The dual effects of the reduced live prey numbers and the restriction on predators’ motion by dead prey lead to a rapid decrease in the rate of killing (which marks a departure from linearity as well).

**Fig 6.**
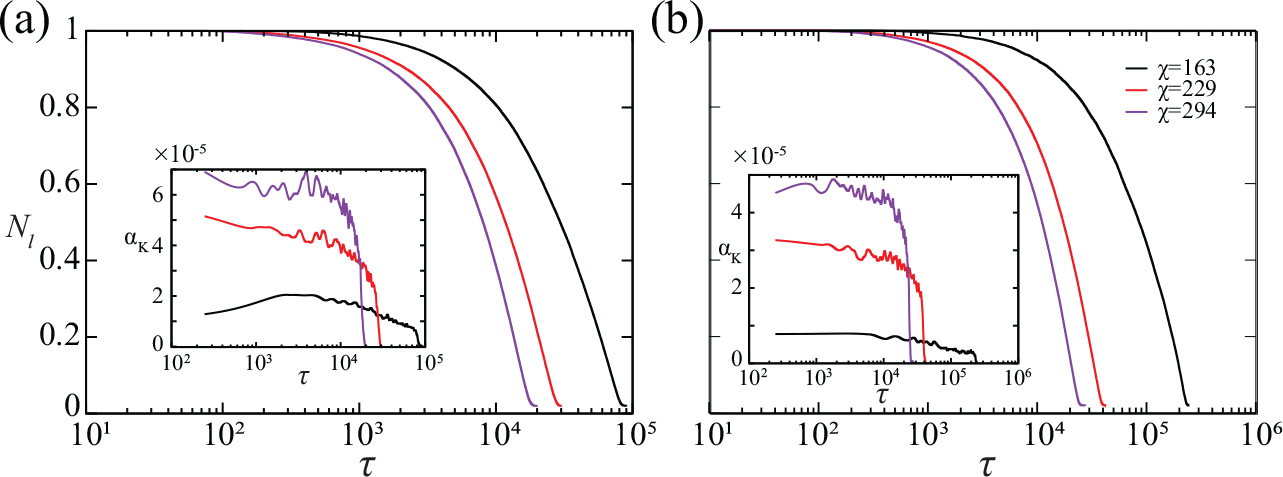
The plots represent the temporal variation of the number of live prey for (a) an all-predator head-on attack and (b) a combination of head-on and split attack. The insets illustrate the change in the rate of killing *α*_*K*_ with time *τ* .

The decay in the number of live prey in the domain presents a general idea of the predators’ hunting prowess. However, the efficacy of the hunting strategy can be defined only by considering all the pursuing scenarios on an encounter basis using the result of the hunt and the time taken for the successful pursuits as metrics. The general notion in predator-prey systems is that the smaller the flock of prey, the easier the hunt [16]. Therefore, it is vital to examine the effect of prey flock size on hunting efficacy; this has been portrayed for different hunting strategies and different degrees of prey coordination in Figs. 7 and 8. In each of the figures, the upper panel portrays the rate of successes and failures for the attacks on different cluster sizes *ϕ*, and the lower panel represents the relation between the elapsed time on successful hunts and the size of the cluster to which the target prey belongs.

**Fig 7.**
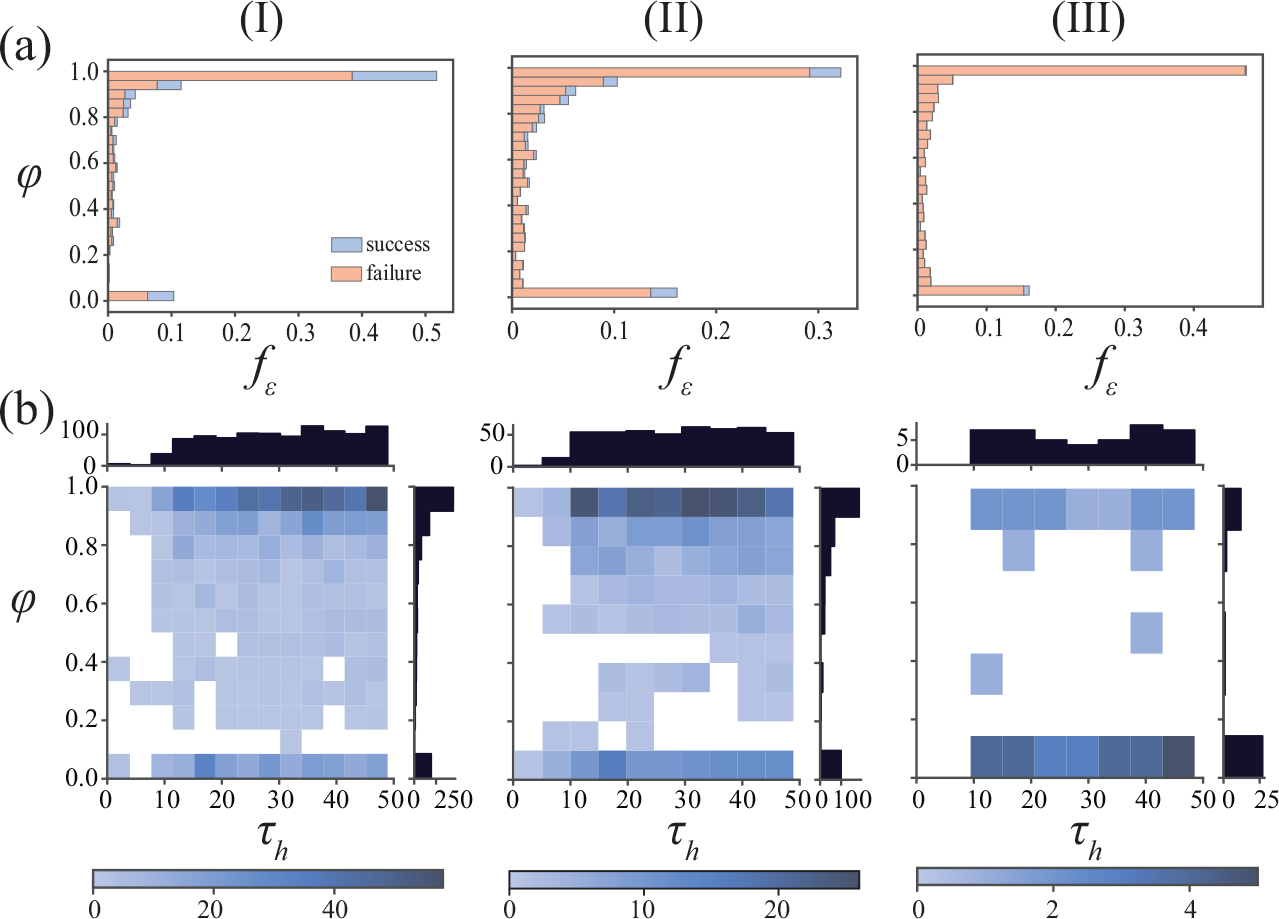
The panel (I) elucidates the success and failure rates of the pursuits against the cluster size *φ*, while the panel (II) portrays the relation between the time required for a successful hunt and the size of the cluster that the target prey belonged to. The columns represent a different set of hunting strategies: (a) represents an all-predator head-on attack, (b) represents a combination of head-on and split attack, and (c) represents an all-predator split attack strategy, respectively. The results are attributed to a lower herding tendency (*χ* = 163).

**Fig 8.**
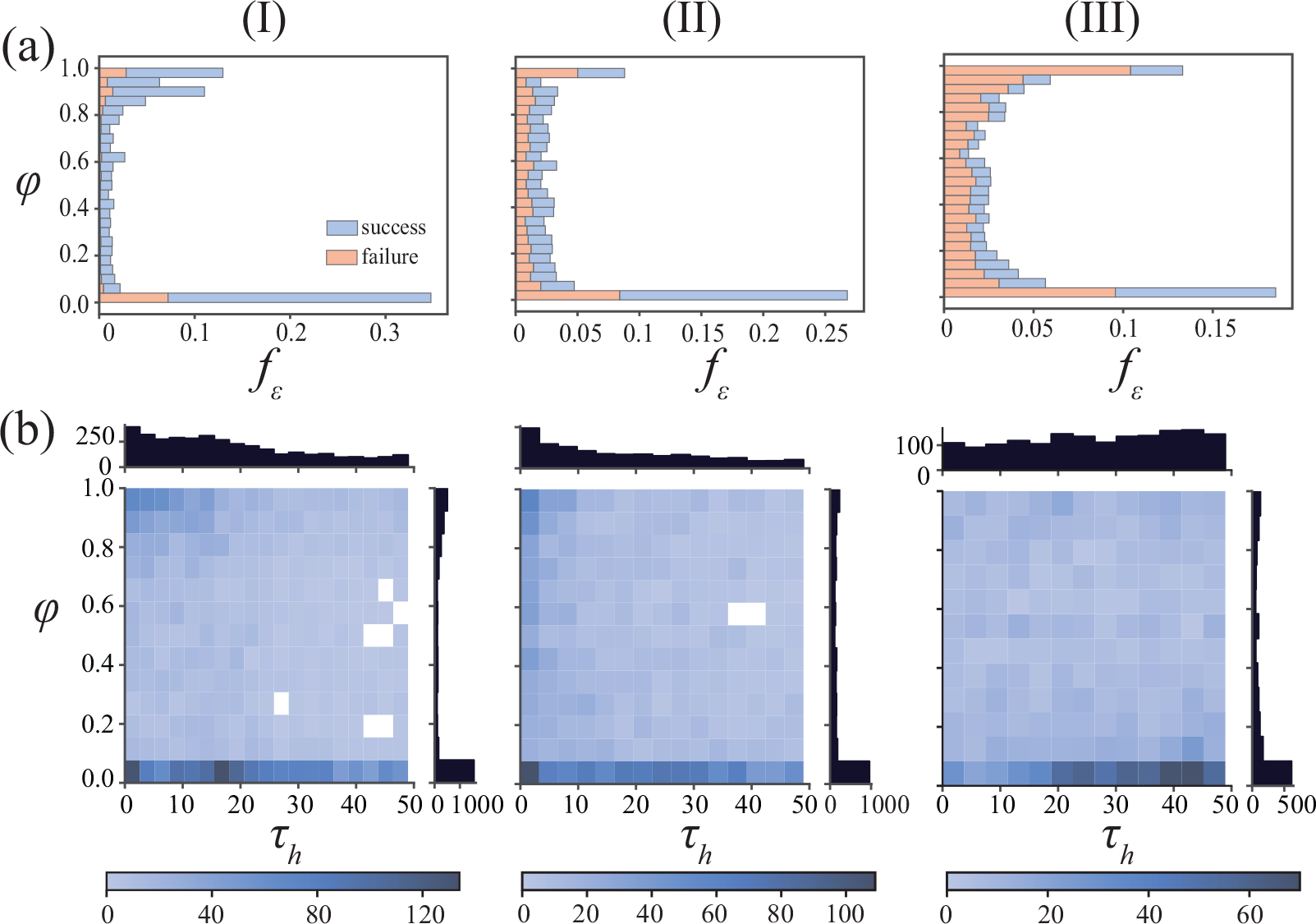
The panel (I) elucidates the success rate of the pursuits against the cluster size *φ*, while the panel (II) portrays the relation between the time required for a successful hunt and the size of the cluster that the target prey belonged to. Each of the columns represents a different set of hunting strategies: (a) represents an all-predator head-on attack, (b) represents a combination of head-on and split attack, and (c) represents an all-predator split attack strategy respectively. The results are attributed to a higher herding tendency (*χ* = 294).

At low herding tendency (*χ* = 163), the majority of the attacks are observed to be focused on larger flock sizes, albeit with a high rate of failure. The purely head-on attack strategy is observed to have the maximum overall success rate at ≈ 28% of all encounters, followed by the combination strategy at ≈ 12% and the split strategy at ≈ 1.1%. The predators following the split strategy fail more often in capturing the target prey successfully, which is understandable as the split strategy is geared towards splitting the flock rather than hunting down prey. Due to this feature of the split strategy, more encounters are recorded involving mid-sized and small-sized clusters in the combination (refer Fig. 7(a-II)) and the split (refer Fig. 7(a-III)) strategies compared to the head-on strategy (see Fig. 7(a-I)). In the existing literature, predators are mostly inclined towards hunting isolated prey [16, 24]. Such behaviour is also observed in the current work, accounting for ≈ 21% of kills in case of head-on attacks, ≈ 25% in case of combination strategy and ≈ 58% in case of split attacks at *χ* = 163. Moreover, in all three strategies, the pursuit time for successful hunts is found to be uniformly distributed from *τ*_*h*_≈ 10 to *τ*_*h*_ ≈ 48, denoting protracted pursuits are just as likely as briefer pursuits.

In case when the prey possesses high herding tendency (*χ* = 294), the head-on strategy still clearly has the highest success rate (≈ 81%) compared to the combination strategy (≈ 60%) and the split strategy (≈ 41%). However, the gap in success rate between the head-on strategy and the split strategy has reduced considerably from around 25 times (at *χ* = 163) to ≈ 2 times (at *χ* = 294). It can also be observed that a significant number of attacks take place at very low cluster sizes, i.e., there is a bias towards isolated prey candidates. Isolated prey account for ≈ 38% of the kills in the case of the head-on strategy compared to ≈ 32% and ≈ 25% in the case of combination and split strategies, respectively. Due to this proclivity towards hunting more isolated prey, the pursuit time in the case of the head-on strategy or the combination strategy is generally on the lower spectrum (see Fig. 8(b-I) and (b-II)), while the split strategy due to its highly fallible nature has a higher average pursuit time (see Fig. 8(b-III)). A similarity with the low herding case is that of the occurrence of mid-sized and small-sized clusters, which is more prominent in the split strategy (see Fig. 8(a-III)). The hunting data at different levels of herding propensity clearly shows that the head-on strategy has the upper hand when the predators do not coordinate (or rather compete) with each other. However, when the prey tends to herd to a greater extent, the rate of success of hunting smaller clusters far exceeds that of larger clusters (by more than twice in most cases), which goes to show the effectiveness of the occasional use of split strategy.

## Discussion

The systematic evaluation of the prey response to repeated attacks by multiple predators allows us to predict prey behaviour under distress. The current work proposes an agent-based model for simulating predator attacks on a prey flock in a periodic domain. The predator’s pursuit strategy in such agent-based models bears an uncanny resemblance to missile guidance systems. In our work, the predators follow the simplest “pure pursuit” approach, where the predator agent dynamically chases the moving prey by accelerating towards it. However, there are more complex schemes, such as deviated pursuit and parallel navigation [50], which can also be used to express predator pursuit. Palmer and Packer classify the predators’ hunting strategy into ambush and coursing [10]. The current work assumes coursing predators persistently pursuing the prey rather than relying on an ambush tactic. In our work, the interactions among the agents are completely based on visual cues, and visual occlusion has not been considered for the sake of simplicity. However, environmental factors such as heterogeneity of the environment and turbidity of the medium can have an effect on the collective dynamics of the prey [58–60]. Some existing literature does, however, consider the finer details of visual acuity while defining models [61, 62]. At the same time, it has been observed that visual cues form the major basis for risk perception in predator-prey systems. The debate on the practicality of aposematism (as a purported defence mechanism) on predation viability advocates the importance of visual cues in prey selection [63, 64]. In the purview of such significant promoters of visual cues, other sensory inputs, such as auditory and haptic, have been neglected in the current work. However, in certain predator-prey pairs, auditory inputs have been found to be the major source of predation risk [65, 66]. The prey evasion, as well, occurs using the simplest possible rules, i.e., accelerating away from the predator. In some animal species, the prey might take an alternate protean approach to evasion [67]. However, a recent study by Szopa-Comley and Ioannou [68], involving robotic prey and natural predators, has challenged the efficiency of the protean approach (i.e., stochastic direction of prey evasion) in the long-term dynamics of such systems. Apart from the prey evasion force or the predating hunting force (i.e., the propulsion force), every agent in the current model is subjected to an attraction force, a friction drag, and an alignment force. The alignment force is the most vital among these forces, as it enables the prey to flock/school instead of swarming. However, at high coordination among the prey, the speed of the prey is observed to diminish, resulting in sluggish motion. A possible cause for this slowdown is the nature of the alignment force used in the model, which causes retardation to any sudden velocity changes in a direction dissimilar from the neighbouring velocity. However, the retardation can be thought of as the cost of alignment for the prey. The current work also assumes non-consuming predators; however, it is observed that consuming predators lead to almost identical long-term behaviour as a non-consuming predator (see S3 Fig).

The problem of interest in the current work is that of the effect of the presence of multiple predators in a predator-prey system. Although the predator-prey dynamics involving a single predator and one or more prey have had their fair share of the limelight, the complexity of the dynamics scales precipitously with even the addition of another predator. The effect of multiple predators is clearly non-additive, as can be seen in S5 Fig. Even discounting the combinatorics of the prey selection strategy, parameters such as the relative angle of attack and synchronisation (phase) dynamics of the attacks play a major role in deciding the efficacy of the hunting process. With the change in the relative angle of attack, a transition from the herding to the splitting regimes is observed. Prey escape manoeuvres such as split-and-join, herd, flash expansion, vacuole, ball, and avoid, some of which have been recreated in the current work, are found to occur in nature widely [6, 56, 69]. The split-and-join behaviour is especially feature-rich and occurs naturally for evading lone predators as well. However, when multiple predators are involved, a slight phase difference (i.e., mismatched) in the attack timings can lead to disastrous consequences for the prey, possibly even to the tune that the escape manoeuvre is rendered useless to the second attack. It is found that when multiple prey flocks join into one, there is a duration when the prey are befuddled with respect to the general direction of movement. A well-timed attack by a predator can lead to higher prey mortality rates. The role of asynchrony is crucial in determining optimal attack patterns in systems with multiple predators, as has been carried out for lone predators in extant literature [24, 36]. As the disturbance in the order caused by every predator attack outlives the attack duration, the frequency of attacks extensively affects the efficacy of prey escape manoeuvres. Theibault et al. [37] examine the improvement in hunting success per predator with successive attacks on a prey aggregate from experimental observations, while Lett et al. [36] define an attraction-alignment-repulsion model and explore the effect of frequency of predator attacks on the flash expansion characteristics of the prey flock. Group hunting tactics are another way for predators to increase their hunting success, as has been reported in African lions [41, 42], African wild dogs [43], and killer whales [44] among others. The current work examines the efficacy of different attack strategies, highlighting the effectiveness of the split strategy in dividing flocks and throwing the prey flock into confusion while depicting the high hunting success rates associated with a head-on attack. When used in tandem in a synchronised fashion, a combination of these attacks can nudge the hunting success rates even higher (as is observed in the works on group hunting).

The model explained in the current work provides us with a framework to explain the ephemeral and long-term behavioural dynamics of a prey flock when attacked by multiple competing predators. A subsequent course of action would be to understand the mechanism of group hunting, which remains ambiguous despite the numerous observations in the wild. The major advantage of using such an agent-based model is the ease of fitting the model to different predator-prey pairs based on their physical attributes and available experimental data. However, as mentioned earlier, there are certain assumptions considered while formulating the model, which leaves room for improvement. Lastly, the effect of environmental factors such as turbulence, obstacles on the marine floor, ocean currents and turbidity of the medium, neglected currently for the ease of modelling, could also be scrutinised.

## Supporting information

**S1 Appendix Scaling analysis of prey speed with coordination parameter**.

(PDF)

**S2 Appendix Determining the nature of the decay in live prey numbers with time**.

(PDF)

**S1 Fig. The process of predation has been explained in the form of a flowchart**. The predation process takes place in four steps: a) detection stage, b) target selection, c) pursuit stage, and d) satisfaction/refocus stage. In the detection stage, the predator searches its visual neighbourhood (detection zone sans the blind spot region) for live prey. In the next step, the predator selects a target prey based on the hunting strategy (head-on or split) it follows. Once a target prey is selected, the predator starts stalking and pursuing the target for a certain maximum hunting time *τ*_*H*_. If the predator is able to kill the target prey (i.e., if the target prey moves into the kill zone of the predator), it enters a satisfaction stage (*τ*_*S*_), where it cruises around without hunting. On the contrary, if the predator is unsuccessful in killing the target prey within the stipulated time period, it enters a refocus stage (i.e., it takes time *τ*_*R*_ to collect its wits). After the satisfaction time *τ*_*S*_ or refocus time *τ*_*R*_ has passed, the predator actively starts detecting potential target prey again. The above predation process has been reported by Romenskyy et al. [16] and used a priori by Demsar et al. [24]. (PDF)

**S2 Fig. Asynchrony of predators**. The model doesn’t specify any rules for synchronisation among predators, i.e., there is no interaction rule between the predators apart from the volume exclusion. The figure displays the temporal variation of the state of the predators. A predator can have three distinct states: a pursuit state (denoted here by 0), a satisfaction state (denoted by 1) and a refocus state (denoted by − 1). The time series of the two predators’ states show moments of asynchrony as well as coincidental synchrony.

(PDF)

**S3 Fig. Comparison of a consuming and a non-consuming predator**. The temporal variation of the number of live prey *N*_*l*_ has been compared for a consuming predator (prey vanishes, once dead) and a non-consuming predator (dead prey act as obstacles, instead of vanishing) for different degrees of coordination *χ*. The long-term dynamics of the two systems are almost identical. (Note: Both the predators follow the head-on attack strategy in the illustrated case; *χ* = 294.)

(PDF)

**S4 Fig. Comparison of deterministic and stochastic prey selection for combination hunt strategy**. In the case of deterministic prey selection for the combination strategy (column a), one predator is set to follow head-on and the other is set to follow split strategies throughout the simulation time. On the other hand, in the case of stochastic prey selection (column b), it is equally probable for each predator to select between the head-on and the split strategies. The upper row (I) shows the success rate of predator attacks on varying normalised prey flock sizes *φ*, while the lower row (II) illustrates the time required *τ*_*h*_ for mounting the successful attacks on these flock sizes. (Note: These statistics are representative of a high herding tendency (*χ* = 294).) (PDF)

**S5 Fig. Comparison of long-term hunting statistics of a single predator and two predators**. The temporal variation of the fraction of prey alive *N*_*l*_ is illustrated for the cases of a single predator following a head-on hunting approach and two predators following a head-on hunting strategy. The inset shows the ratio (*κ*) of the time taken for a lone predator to hunt a certain fraction of prey agents *N*_*d*_. A solitary predator takes much longer to hunt at large flock sizes compared to a predator duo. The time difference decreases as the flock size starts diminishing. However, the time taken by one predator to hunt a certain number of prey is always found to be more than twice that taken by two predators (i.e., the effect of multiple predators is not additive). (Note: These statistics are representative of a high herding tendency (*χ* = 294).) (PDF)

**S1 Table Parameter space**. Due to the complexity of the model, the parameter space of the model has been explored through preliminary simulations, and the parameter values used in the simulations have been tabulated.

(PDF)

## Acknowledgments

The authors acknowledge the funding received from MHRD, Government of India, under Project No. SB22230157MEPMRF008846 and the funding received as part of the Institution of Eminence scheme of the Ministry of Education, Government of India [Sanction No: SB22231233MEETWO008509].

